# Shikonin induces odontoblastic differentiation of dental pulp stem cells via AKT–mTOR signaling in the presence of CD44

**DOI:** 10.1101/2020.01.27.921015

**Authors:** Kunihiro Kajiura, Naoki Umemura, Emika Ohkoshi, Takahisa Ohta, Nobuo Kondoh, Satoshi Kawano

## Abstract

In our previous study, we demonstrated that hyaluronan induces odontoblastic differentiation of dental pulp stem cells via interactions with CD44. However, it remains unclear whether CD44 expression by dental pulp stem cells is required for odontoblastic differentiation. Therefore, we searched for a compound that induces odontoblastic differentiation of dental pulp stem cells, regardless of the chemical structure and function of hyaluronan, and examined whether CD44 is involved in the induction of odontoblastic differentiation by the compound. Because vitamin K analogues can promote bone formation and tissue calcification, we focused on derivatives of naphthoquinone, the skeleton of vitamin K; we verified whether those compounds could induce odontoblastic differentiation of dental pulp stem cells. We found that dentin sialophosphoprotein, a marker of odontoblasts, was expressed in dental pulp stem cells after treatment with shikonin. The shikonin-induced expression of dentin sialophosphoprotein was inhibited by PI3K, AKT, and mTOR inhibitors. In addition, in dental pulp stem cells transfected with siRNA against CD44, the shikonin-induced expression of dentin sialophosphoprotein was inhibited. Thus, shikonin can stimulate dental pulp stem cells to undergo odontoblastic differentiation through a mechanism involving the AKT–mTOR signaling pathway and CD44. Hyaluronan stimulated dental pulp stem cells to undergo CD44-mediated odontoblastic differentiation in our previous study; the present study indicated that CD44 is necessary for dental pulp stem cells to undergo odontoblastic differentiation. Although expression of CD44 is important for inducing odontoblastic differentiation of dental pulp stem cells, the relationship between the AKT–mTOR signaling pathway and CD44 expression, in the context of shikonin stimulation, has not yet been elucidated. This study suggested that shikonin may be useful for inducing odontoblastic differentiation of dental pulp stem cells, and that it may have clinical applications, including protection of dental pulp.

## Introduction

In our previous study, we demonstrated that hyaluronan, a large molecule, induces odontoblastic differentiation of dental pulp stem cells (DPSCs) via interactions with CD44 [1]. However, the underlying cellular signaling mechanism has not yet been elucidated, and it has been unclear whether small molecules can induce odontoblastic differentiation of DPSCs. Therefore, we explored whether small molecules could induce odontoblastic differentiation of DPSCs and attempted to identify the differentiation induction mechanism. In addition, we assessed whether CD44 was essential for this mechanism.

In the field of DPSC research, several research groups have shown that collagenase-treated cells isolated from human dental pulp showed more rapid proliferation, compared with hematopoietic stem cells, and a similar degree of multipotency [2,3]. Subsequently, various groups have described possible DPSCs. There have been reports of the induction of odontogenic differentiation from DPSCs in vitro [4–6]. Moreover, human DPSCs can differentiate into hepatocytes [7], and the differentiation of DPSCs into pancreatic β cells can be induced by specific culture conditions [8]. Some in vivo experiments in rats have demonstrated that DPSC transplantation can regenerate corneal epithelium [9], myocardium [10], spinal cord [11,12], muscle (in a model of muscular dystrophy) [13], and craniofacial bone (using collagen gel as a scaffold) [14].

Notably, there are reports that cells expressing CD44, a marker of DPSCs, are present in dental pulp tissue around the tooth root, and that these CD44-positive cells are involved in mineralization [15,16]. Furthermore, our previous study revealed that DPSCs express high levels of CD44; it also showed that hyaluronan, a ligand of CD44, can induce odontoblastic differentiation of DPSCs. These findings suggest that dental pulp cells, including DPSCs, differentiate into odontoblasts through a CD44-mediated mechanism and participate in calcification.

Although the intracellular signaling of DPSCs stimulated with hyaluronan involves activation of various signals (e.g., MAPK and AKT) in a CD44-mediated manner, the causal relationship between odontoblastic differentiation of DPSCs and these activating signals is not clear. To the best of our knowledge, there have been few reports of inducers or factors (e.g., hyaluronan) that efficiently induce odontoblastic differentiation of DPSCs. Furthermore, it remains unclear whether CD44 is required to induce odontoblastic differentiation of DPSCs. Notably, menatetrenone (a vitamin K_2_ analogue) is used clinically as an osteoporosis treatment and hemostatic agent; it is known to be involved in bone formation, tissue calcification, and blood coagulation [17,18]. However, the clinical significance of menatetrenone in the dental field has not been fully investigated. In this study, we focused on vitamin K analogues containing naphthoquinone derivatives, to determine whether these analogues can induce odontoblastic differentiation of DPSCs. Then, we explored small molecules involved in inducing odontoblastic differentiation of DPSCs; the findings enabled us to elucidate whether the intracellular signaling of new compounds that induce odontoblastic differentiation of DPSCs involve CD44-mediated signaling, similar to that observed upon stimulation with hyaluronan.

## Materials and methods

### Reagents and cell culture

Vitamin K_1_, 2-methyl-1,4-naphthoquinone (Vitamin K_3_), vitamin K_4_, 1,4-benzoquinone, emodin, and aloe emodin were purchased from Tokyo Chemical Industry Co., Ltd. (Tokyo, Japan). Vitamin K_2_ and shikonin standard sample (99%) were purchased from Fujifilm Wako Pure Chemical Industries (Osaka, Japan). These reagents were each dissolved in dimethyl sulfoxide as stock solutions (100 mM final concentration) and stored in the dark at −20℃. Human DPSCs were obtained from Lonza Walkersville, Inc. (Walkersville, MD, USA). Cultures were maintained in Dental Pulp Stem Cell Growth Media (Lonza) at 37°C in a humidified atmosphere containing 5% CO_2_.

### Immunoblot analyses

Whole cell extracts were obtained using a lysis buffer (10× RIPA buffer [Cell Signaling Technology, Beverly, MA, USA], with 1 mM phenylmethylsulfonyl fluoride and 1× protease inhibitor). Ten micrograms of protein extracts were separated by electrophoresis in an 8% sodium dodecyl sulfate-polyacrylamide gel and transferred to a polyvinylidene difluoride membrane. Membranes were probed with primary antibodies to CD44 (mouse monoclonal, catalog #5640), phospho-AKT (rabbit monoclonal, catalog #4060), AKT (rabbit monoclonal, catalog #4691), phospho-mTOR (Ser2448) (rabbit monoclonal, catalog #5536), mTOR (rabbit monoclonal, catalog #4691) (all from Cell Signaling Technology; dilution 1:1,000); DSPP (mouse monoclonal, catalog #sc-73632) (Santa Cruz Biotechnology, Inc., Dallas, TX, USA; dilution 1:200); and beta-actin (catalog #A5441) (Sigma-Aldrich, St. Louis, MO, USA; dilution 1:10,000). Signals were revealed using peroxidase-conjugated secondary antibodies; bands were visualized by chemiluminescence (Clarify Western ECL substrate; Bio-Rad, Hercules, CA, USA). The blots and images were developed with the ImageQuant^™^ LAS500 Imaging System (GE Healthcare Bio-Science AB, Uppsala, Sweden).

### Cell viability assay

DPSCs were seeded in 96-well plates at a density of 1×10^3^ cells/well and allowed to adhere for 24 h. Cell were treated with each compound for 48 h. Cell viability was assessed by addition of 5 μl of 3-(4,5-dimethylthiazol-2-yl)-2,5-diphenyltetrazolium bromide (i.e., MTT) using a Cell Proliferation Kit I (Roche Diagnostics, Mannheim, Germany), in accordance with the manufacturer’s instructions. The numbers of viable cells were assessed by measuring the absorbance of the resulting formazan crystals at 595 nm with a TECAN SpectraFluor and XFluor4 software (Tecan Japan Co., Ltd., Kawasaki, Japan). All data are presented as the means ± standard deviations of at least three independent experiments.

### Signal blocking assay

LY294002 (phosphoinositide 3-kinase [PI3K] inhibitor), GSK690693 (AKT inhibitor), and rapamycin (mTOR inhibitor) were purchased from Selleck Chem (Tokyo, Japan). DPSCs were pretreated with inhibitors for 30 min before stimulation with shikonin 0.5 μM. After 30 min of stimulation with shikonin, cells were harvested to investigate the inhibition of several signaling pathways. After 24 h of stimulation with shikonin, cells were harvested to assay the expression of the odontoblastic differentiation marker, dentin sialophosphoprotein (DSPP), by immunoblotting.

### CD44 knockdown by siRNA transfection

Small interfering RNAs (siRNAs) specifically targeting human CD44 (siRNA IDs s2681 and s2682) were obtained from Life Technologies (Carlsbad, CA, USA). DPSCs (1×10^5^ per well) were plated in six-well plates and transfected with CD44 siRNA for 24 h in antibiotic-free media using siRNA Lipofectamine RNAiMAX and OPTI-MEM I reduced serum medium (Invitrogen, Carlsbad, CA, USA), in accordance with the siRNA manufacturer’s protocol. After 24 h of transfection, the cells were used for experiments.

## Results

### Among vitamin K analogues, shikonin induces DPSCs to odontoblastic differentiation

First, we examined whether naphthoquinone derivatives, which constitute the skeleton of vitamin K, could induce odontoblastic differentiation of DPSCs. We found that, among the vitamin K analogues we examined, shikonin prominently induced expression of DSPP, a marker of odontoblastic differentiation (Fig 1). This result suggested that shikonin could induce differentiation of DPSCs into odontoblasts.

**Fig 1.**
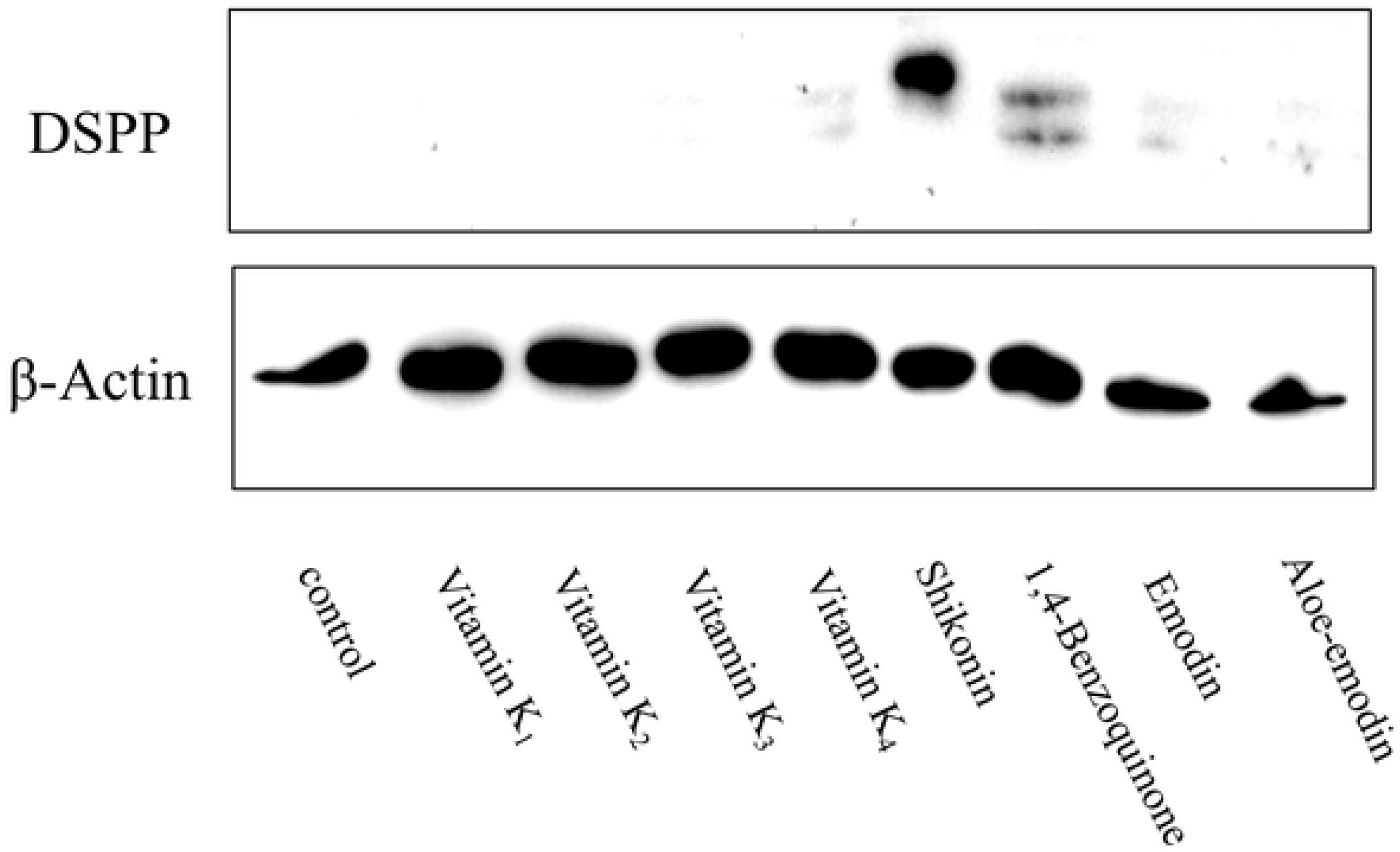
Vitamin K Compounds that Induce Expression of DSPP in DPSCs. The presence or absence of DSPP expression after stimulation with vitamin K compounds was verified by immunoblotting. DPSCs were separately treated with 1 μM of each compound (vitamin K_1_, K_2_, K_3_, K_4_, shikonin, 1,4-Benzoquinone, emodin, and aloe emodin) for 24 h.

### Shikonin induces odontoblastic differentiation of DPSCs, whereas its enantiomer does not

Next, we examined the 50% inhibitory concentrations of shikonin and its enantiomer, alkannin, against DPSCs (Fig 2A). The 50% inhibitory concentrations did not significantly differ between shikonin (4.48 ± 0.51 μM) and alkannin (5.08 ± 1.21 μM) (Table 1). Furthermore, DSPP was expressed by DPSCs stimulated with shikonin concentrations greater than 0.1 μM (Fig 2B). Conversely, stimulation with alkannin did not induce expression of DSPP by DPSCs (Fig 2C). These results suggested that shikonin could induce differentiation of DPSCs to odontoblasts in an enantioselective manner.

Table.1
| 50% inhibitory concentration values (μM) in DPSCs |  |
| --- | --- |
| Compound | IC<sub>50</sub> (μM) |
| Shikonin | 4.48 ± 0.51 |
| Alkannin | 5.08 ± 1.21 |

**Fig 2.**
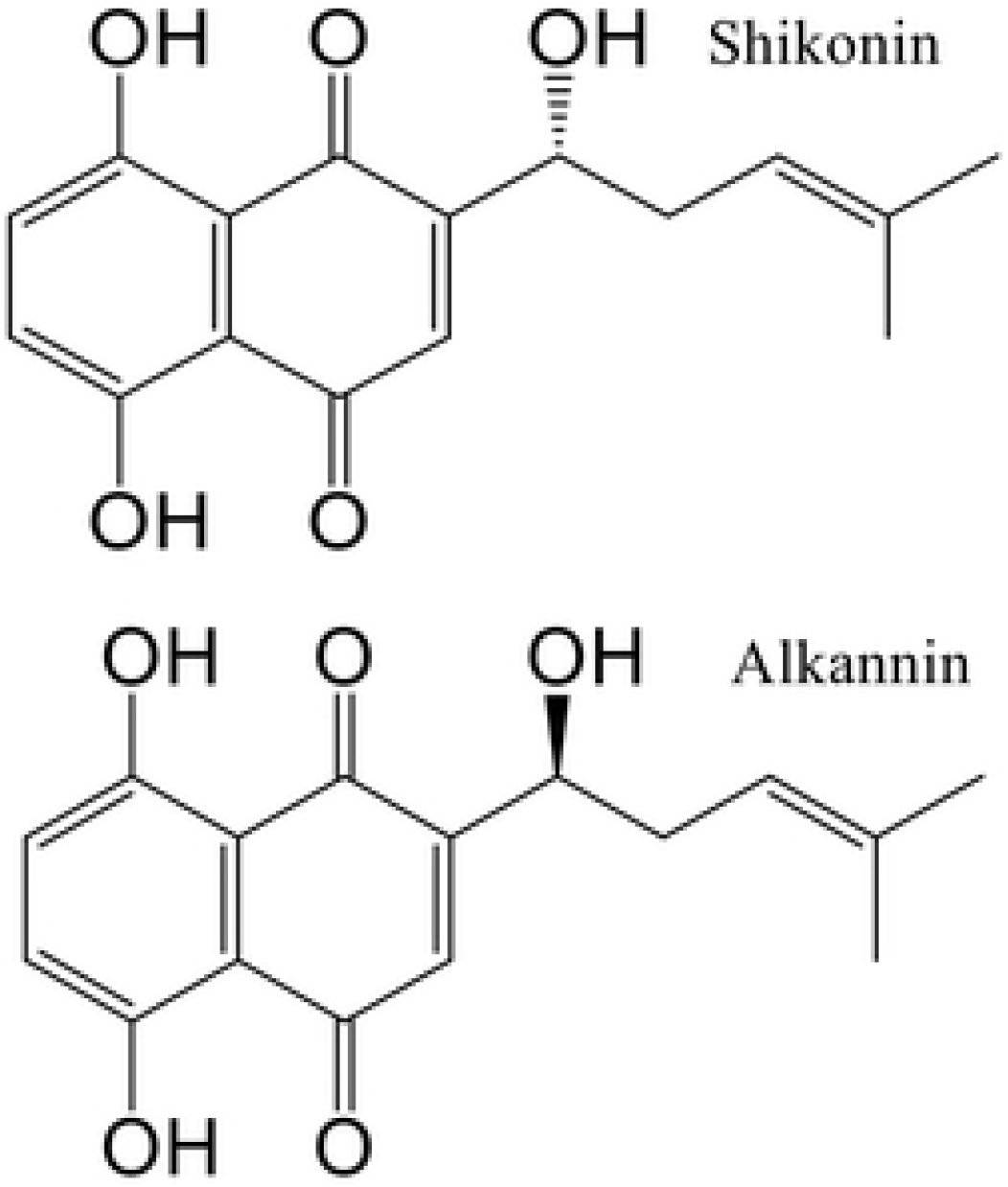
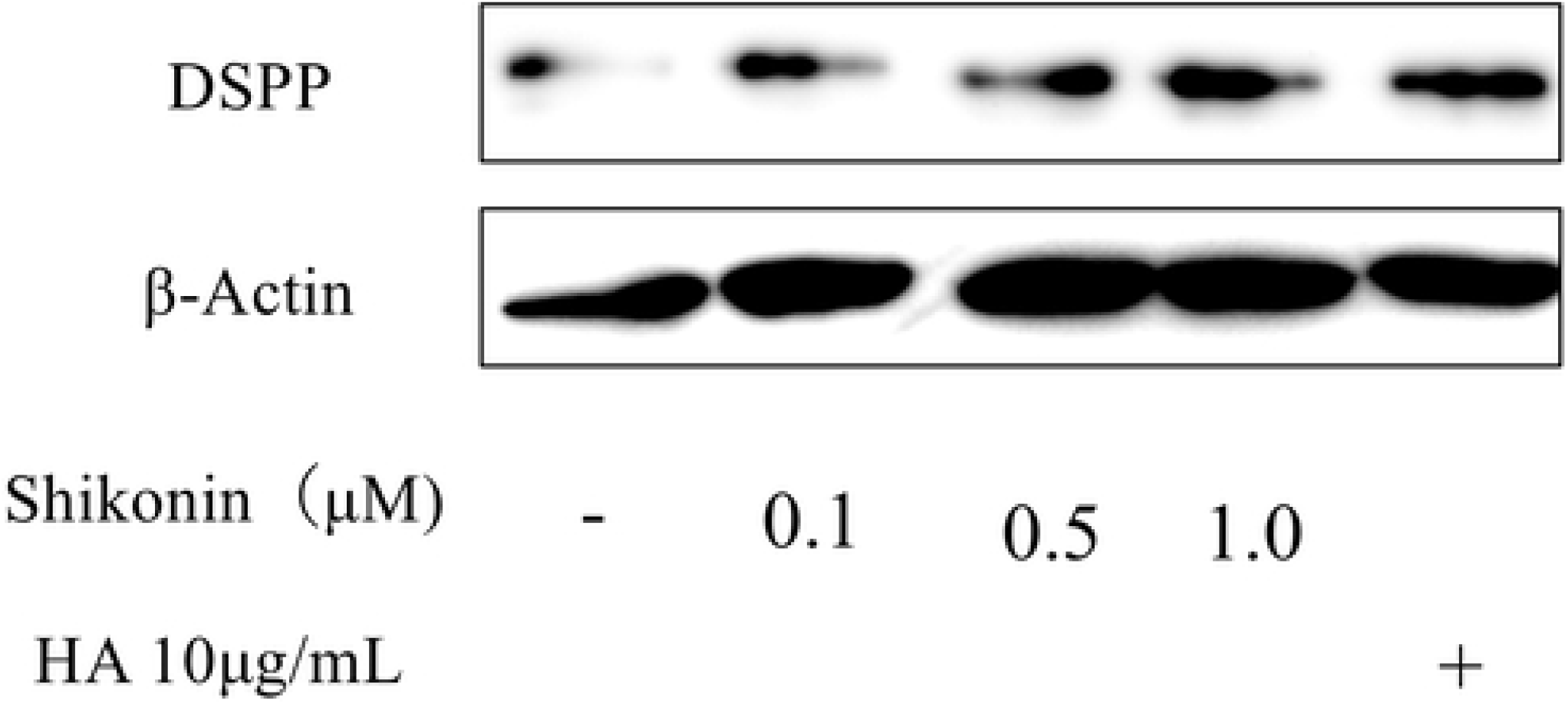
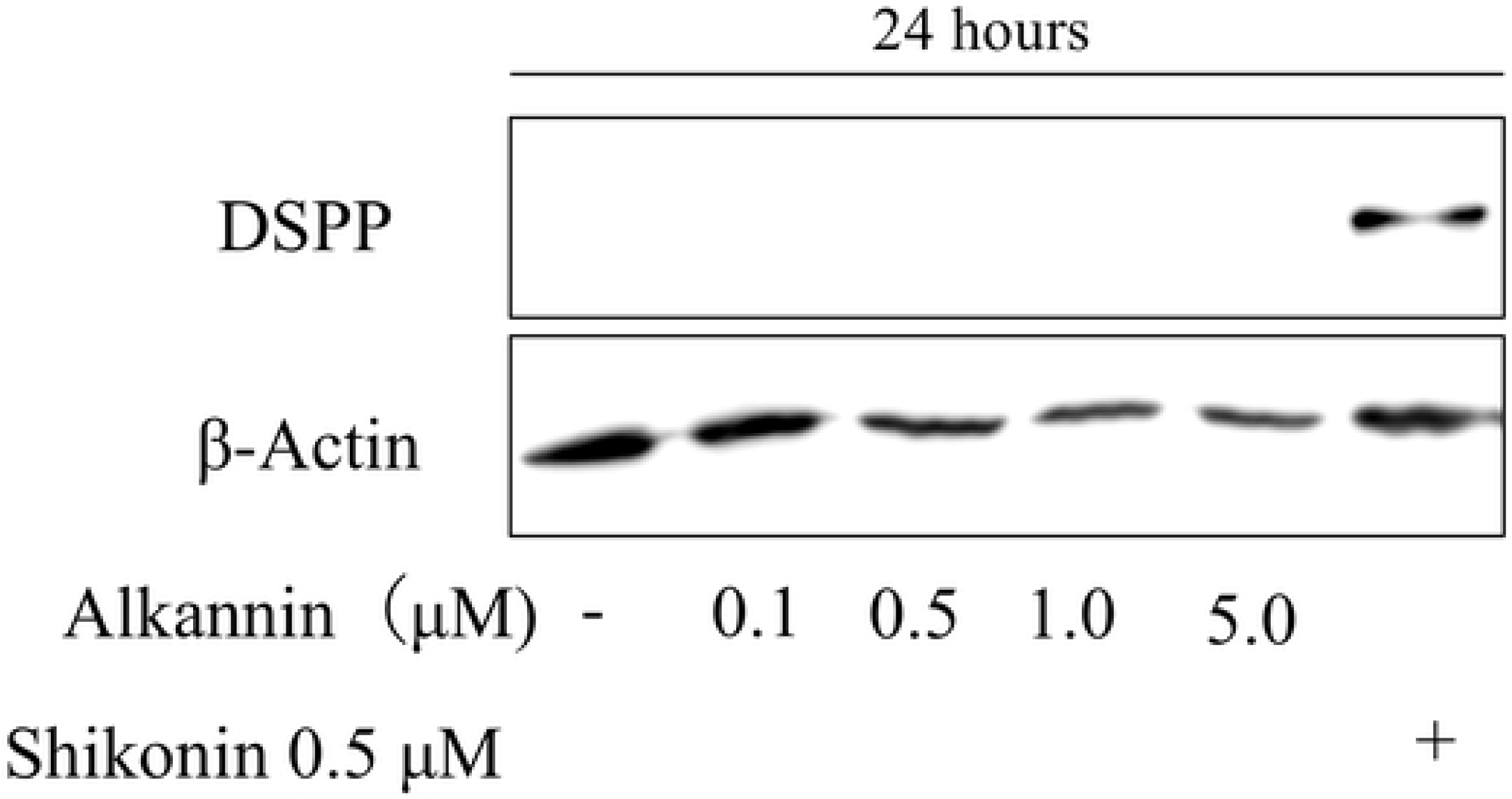
Shikonin Induces Expression of DSPP in DPSCs. The chemical structures of shikonin and its optical isomer, alkannin (A). Induction of the expression of DSPP by shikonin at concentrations ranging from 0.1 μM to 1.0 μM (B). Induction of the expression of DSPP by alkannin at concentrations ranging from 0.1 μM to 5.0 μM (C).

### Shikonin induces odontoblastic differentiation of DPSCs via AKT–mTOR signaling

We investigated the shikonin-related intracellular signal transduction that could induce odontoblastic differentiation of DPSCs. AKT phosphorylation peaked at 30 min after shikonin treatment (Fig 3A); mTOR phosphorylation was also observed between 30 and 90 min after shikonin treatment (Fig 3B). Based on these results, we hypothesized that AKT and mTOR play important roles in the mechanism by which shikonin induces odontoblastic differentiation of DPSCs. Thus, we investigated whether DSPP expression was inhibited upon administration of inhibitors for these proteins. When the phosphorylation of AKT was suppressed by treatment with the PI3K inhibitor, Ly294002, the expression of DSPP was suppressed (Fig 3C, D). In addition, when AKT was inhibited with the pan-AKT kinase inhibitor, GSK690693, the expression of DSPP was suppressed (Fig 3E, F). Furthermore, when phosphorylation of mTOR was inhibited by the mTOR inhibitor, rapamycin, the expression of DSPP was suppressed (Fig 3G, H). These results indicated that shikonin could induce odontoblastic differentiation of DPSCs via AKT–mTOR signaling.

**Fig 3.**
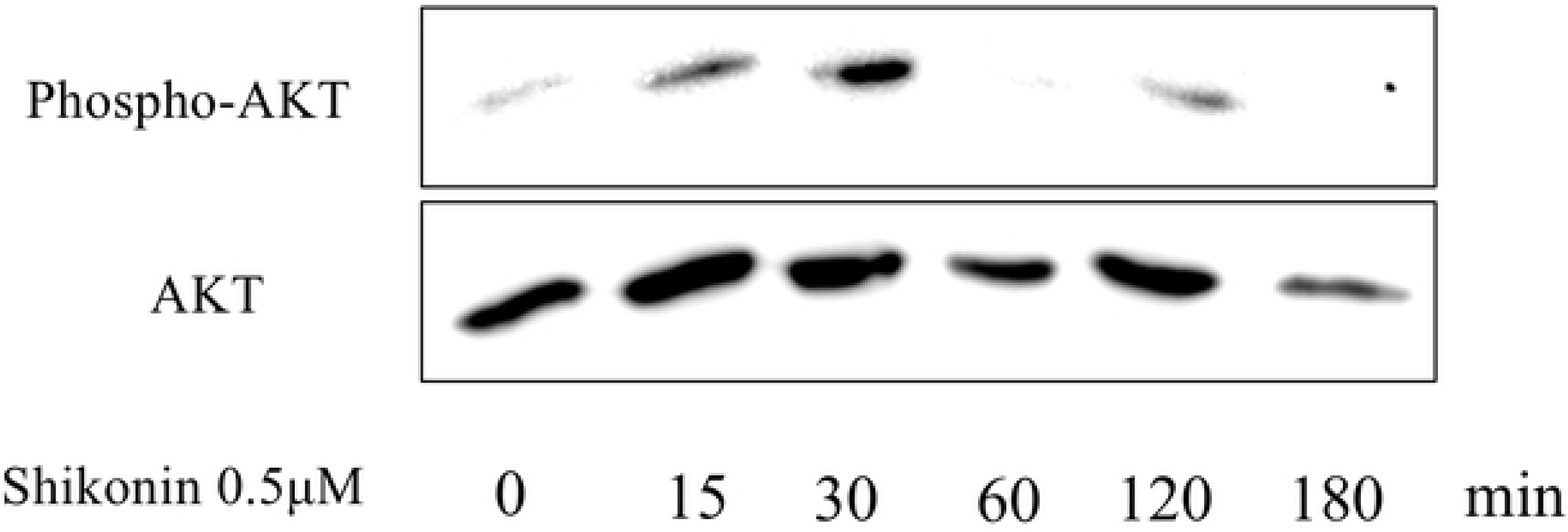
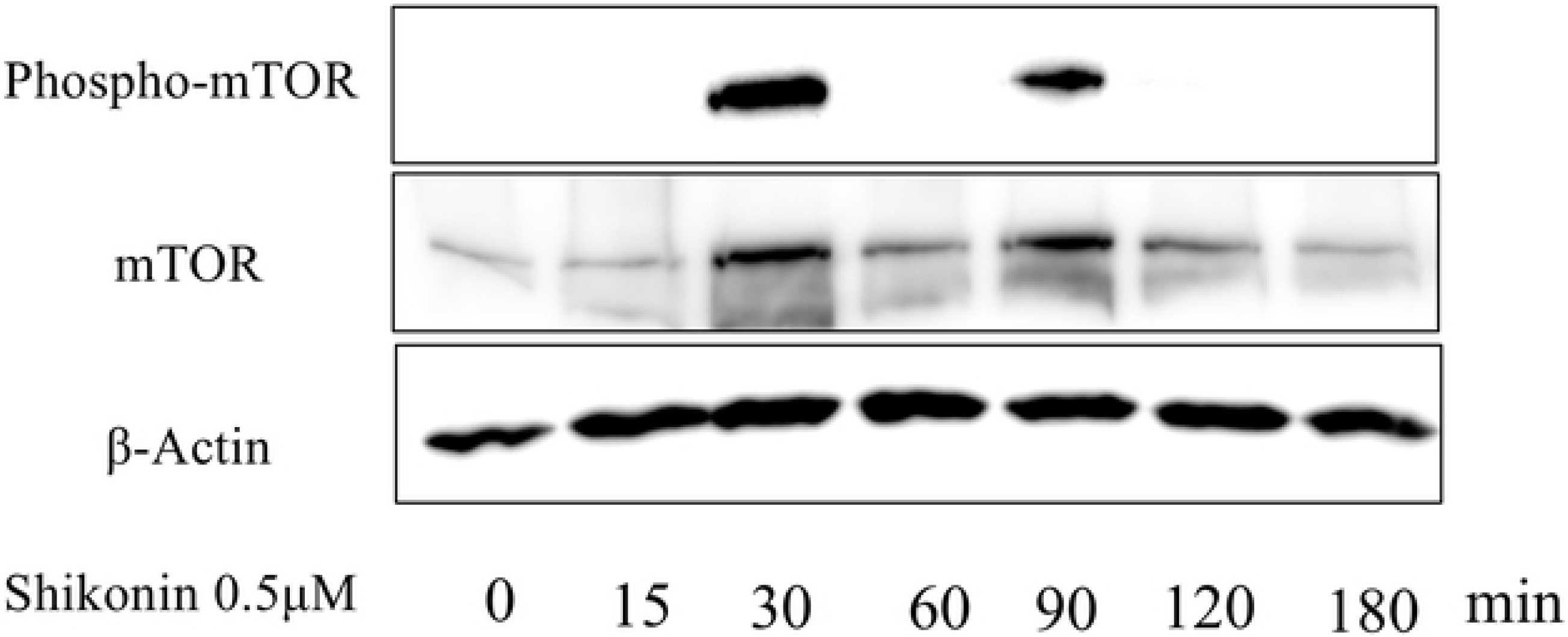
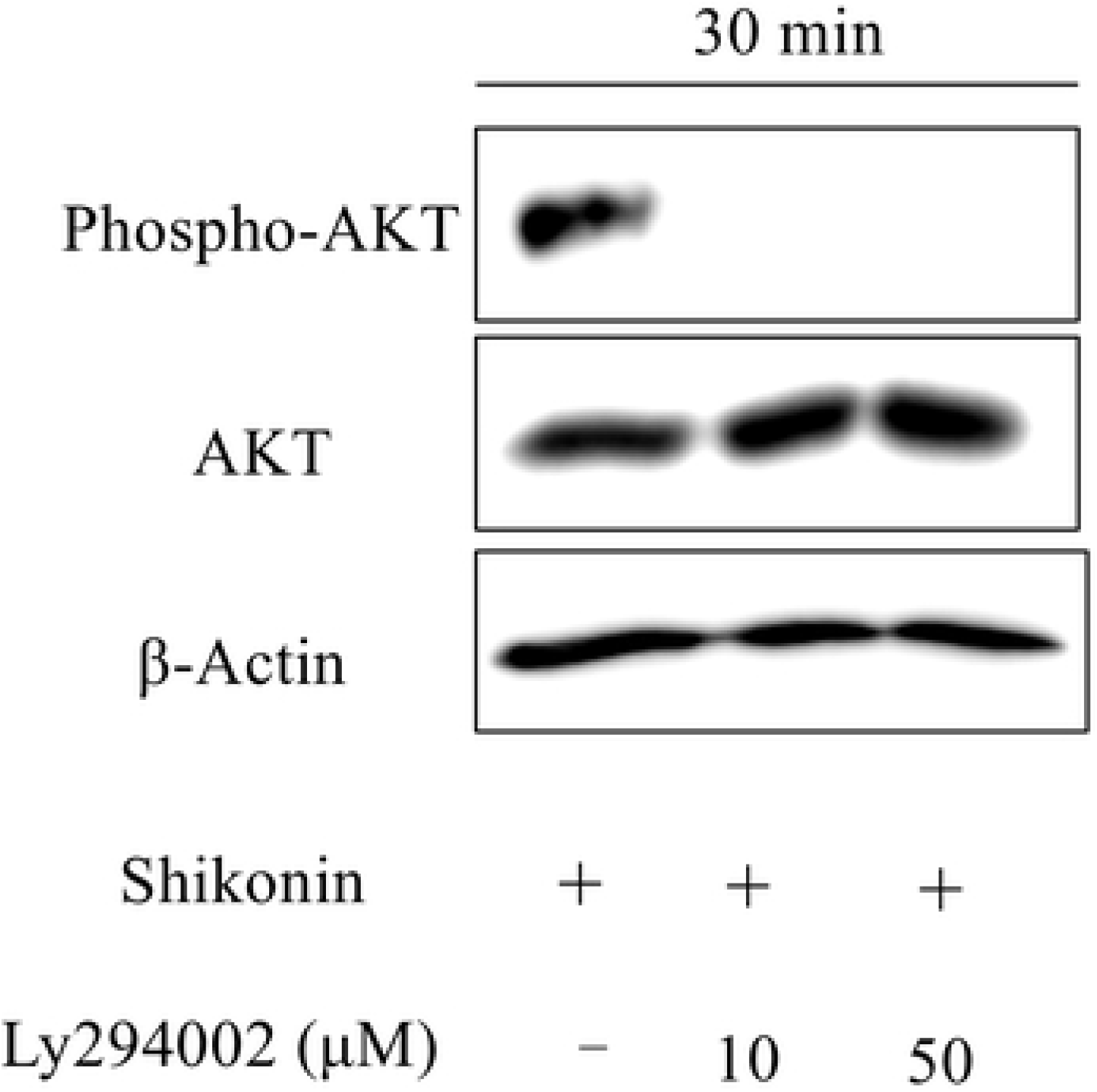
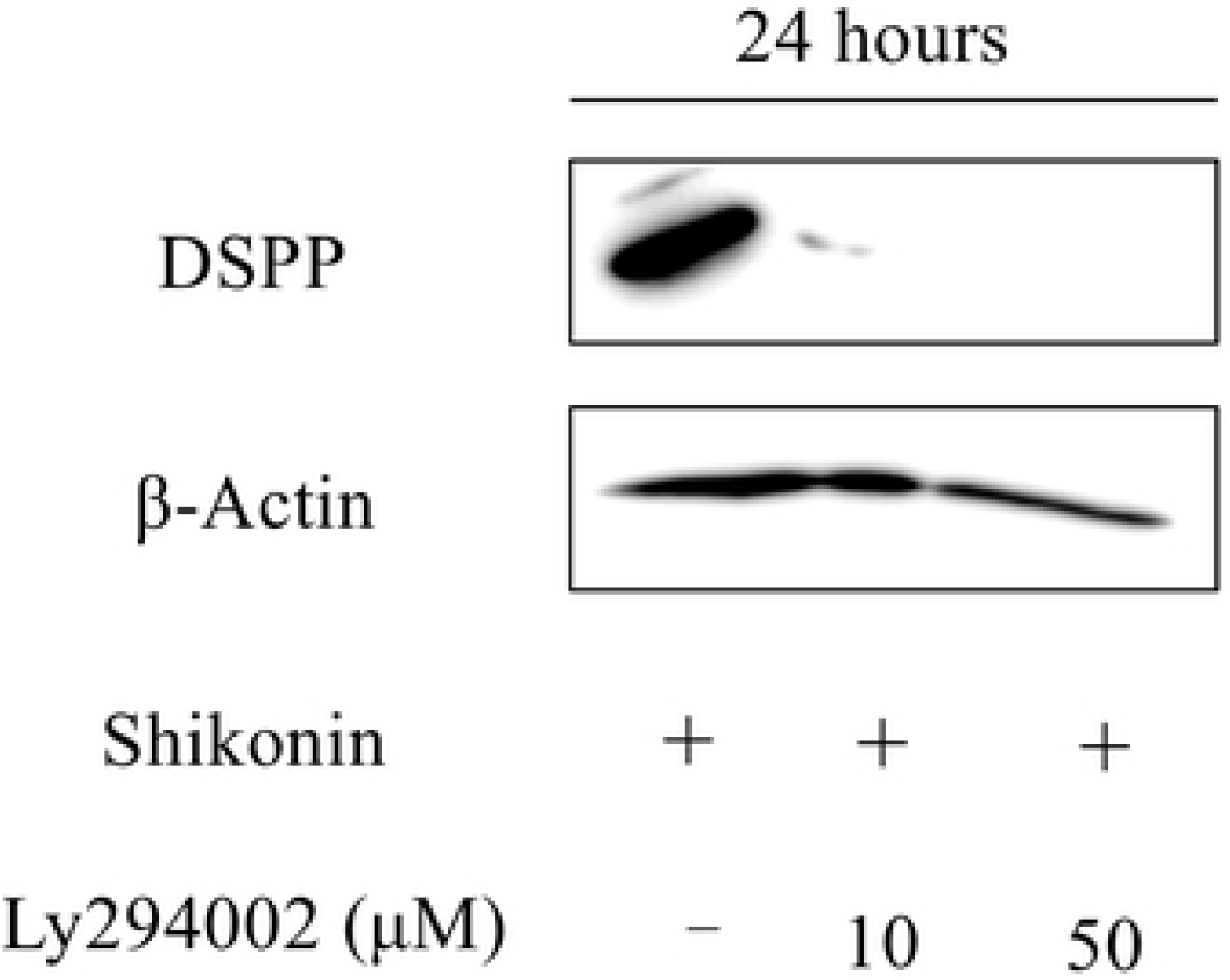
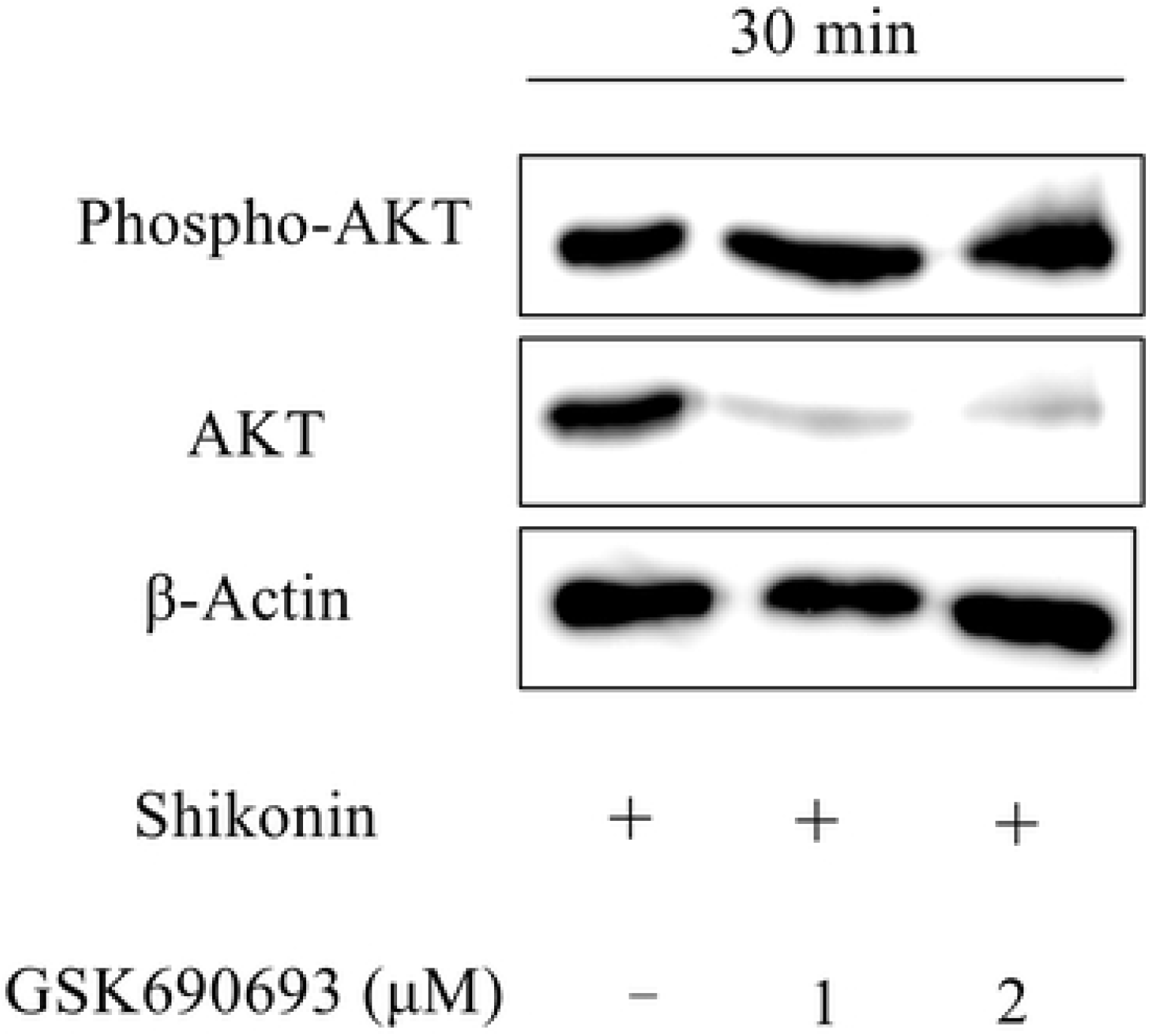
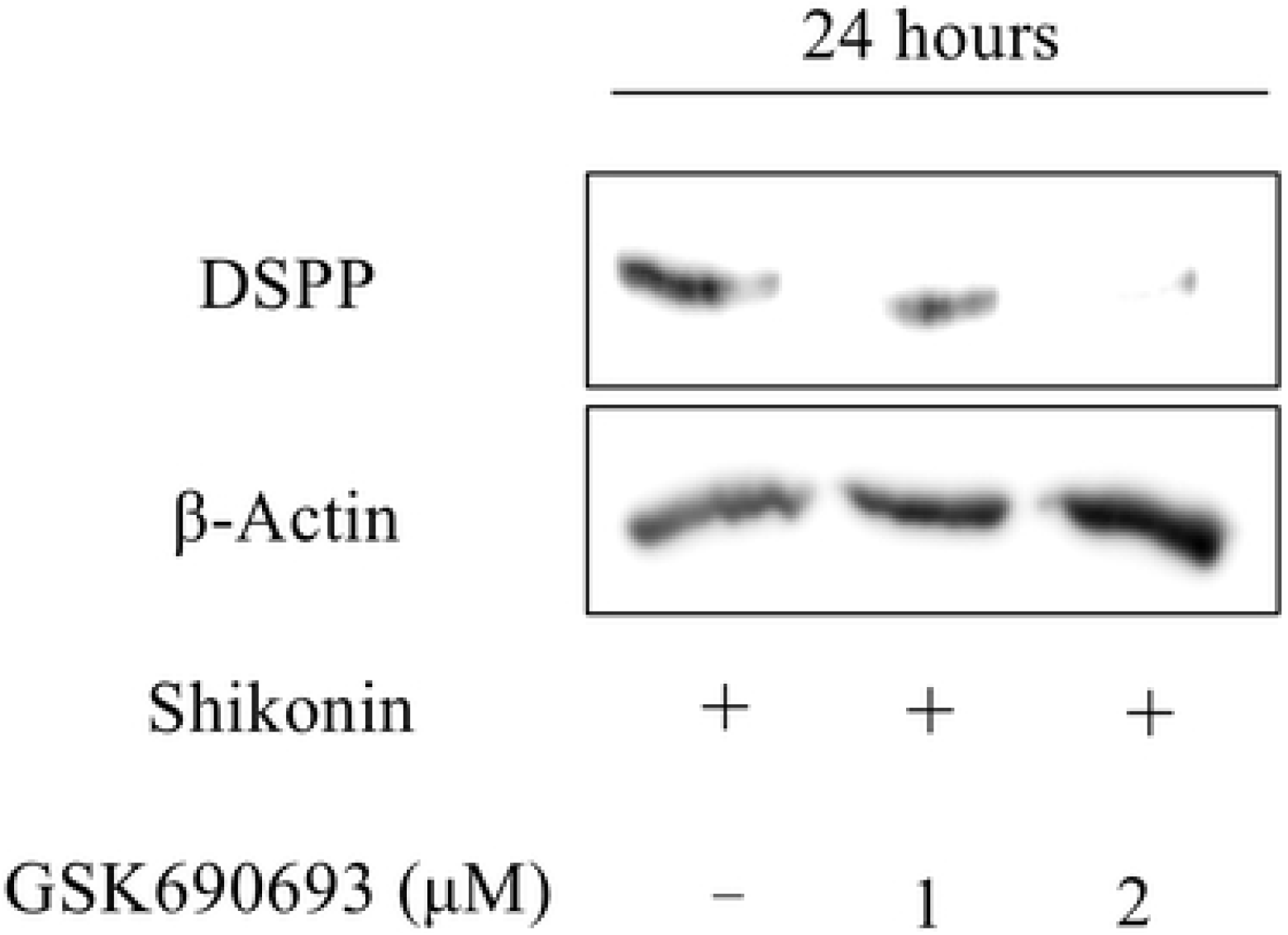
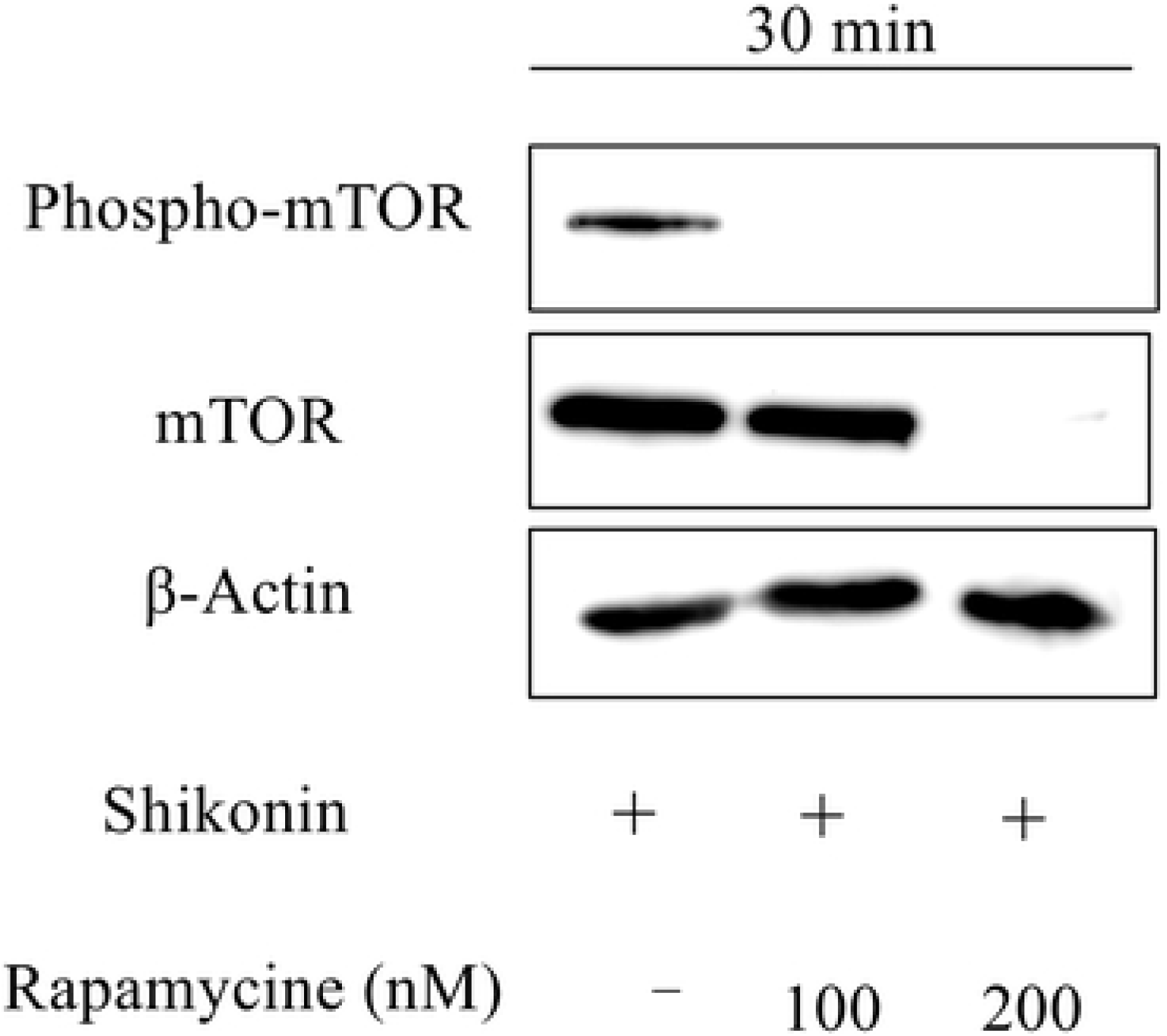
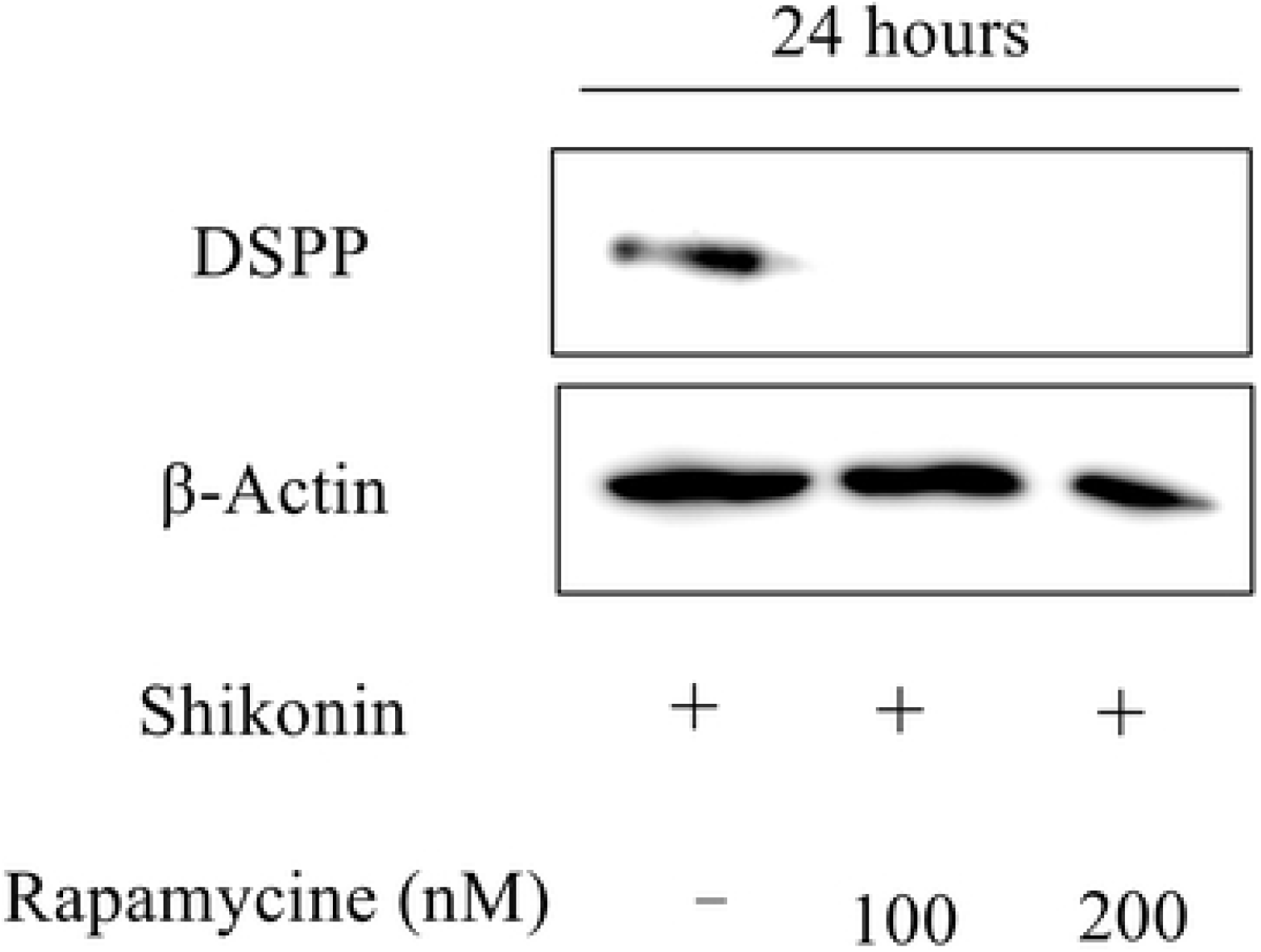
Shikonin-induced Expression of DSPP is Mediated by the AKT–mTOR Signaling Pathway. Phospho-AKT expression levels over time, from 15 min to 180 min, during treatment with 0.5 μM shikonin (A). Phospho-mTOR and mTOR expression levels over time, from 15 min to 180 min, during treatment with 0.5 μM shikonin (B). Phospho-AKT expression levels after co-treatment with 0.5 μM shikonin and LY294002 for 30 min (C). DSPP expression levels after treatment with 0.5 μM shikonin for 24 h, following inhibition of phospho-AKT by treatment with LY294002 (D). AKT expression levels after co-treatment with 0.5 μM shikonin and GSK690693 for 30 min (E). DSPP expression levels after treatment with 0.5 μM shikonin for 24 h, following inhibition of AKT by treatment with GSK690693 (F). mTOR expression levels after co-treatment with 0.5 μM shikonin and rapamycin for 30 min (G). DSPP expression levels after treatment with 0.5 μM shikonin for 24 h, following inhibition of mTOR by treatment with rapamycin (H).

### Shikonin induces CD44-mediated odontoblastic differentiation of DPSCs

To determine whether shikonin could induce CD44-mediated odontoblastic differentiation of DPSCs, we knocked down CD44 expression by using siRNA transfection. siRNA knockdown of CD44 suppressed odontoblastic differentiation of DPSCs, despite shikonin treatment (Fig 4A). To investigate the relationships between shikonin and CD44 with respect to the odontoblastic differentiation of DPSCs via AKT– mTOR signaling, we examined changes in AKT–mTOR signaling after shikonin treatment in DPSCs that had been treated with CD44 siRNA. When CD44 was inhibited, both AKT and mTOR signaling pathways were induced by shikonin treatment (Fig 4B). This result indicated in the absence of CD44, shikonin does not induce odontoblastic differentiation of DPSCs, despite its ability to induce AKT–mTOR intracellular signaling.

**Fig 4.**
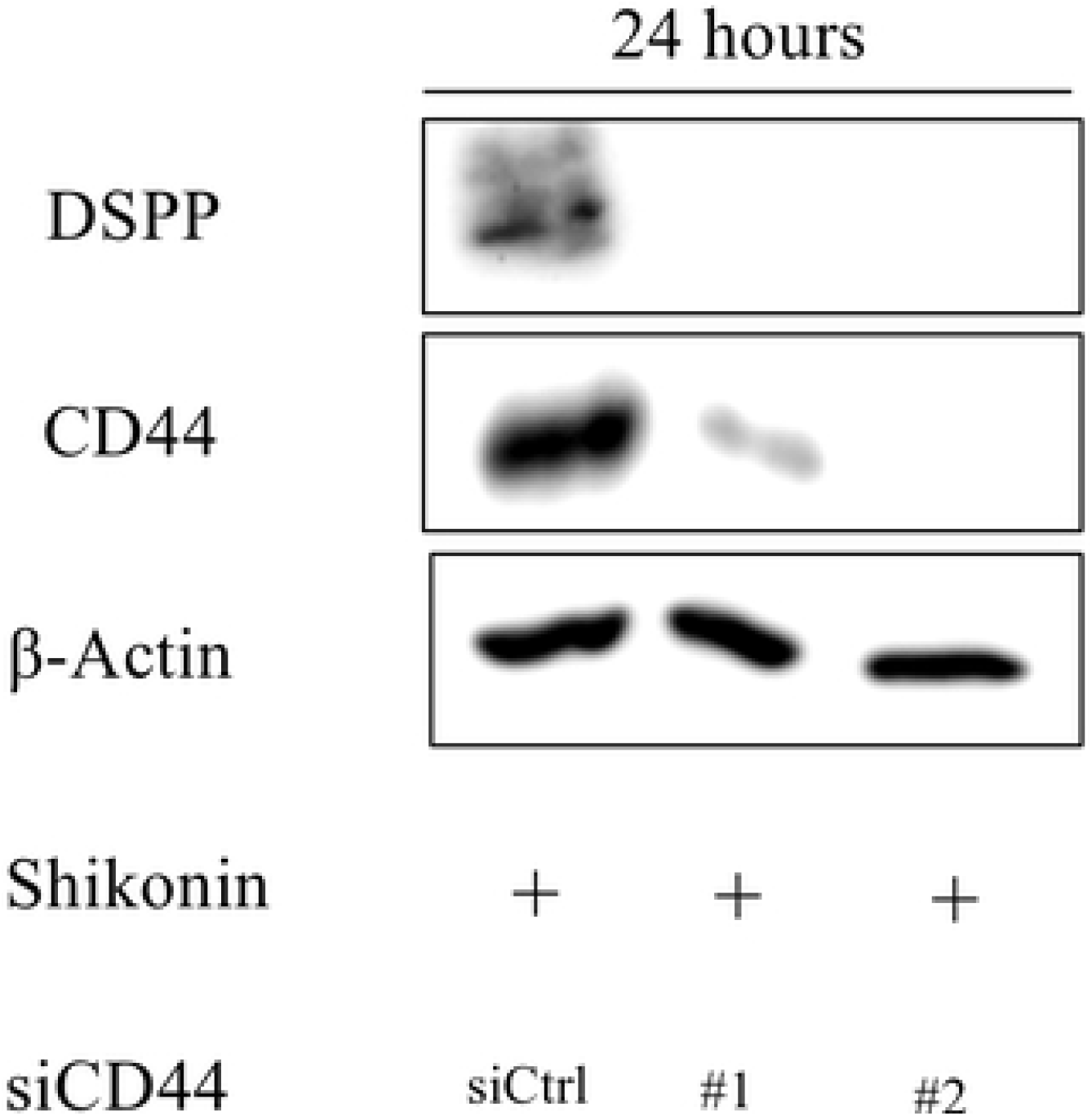
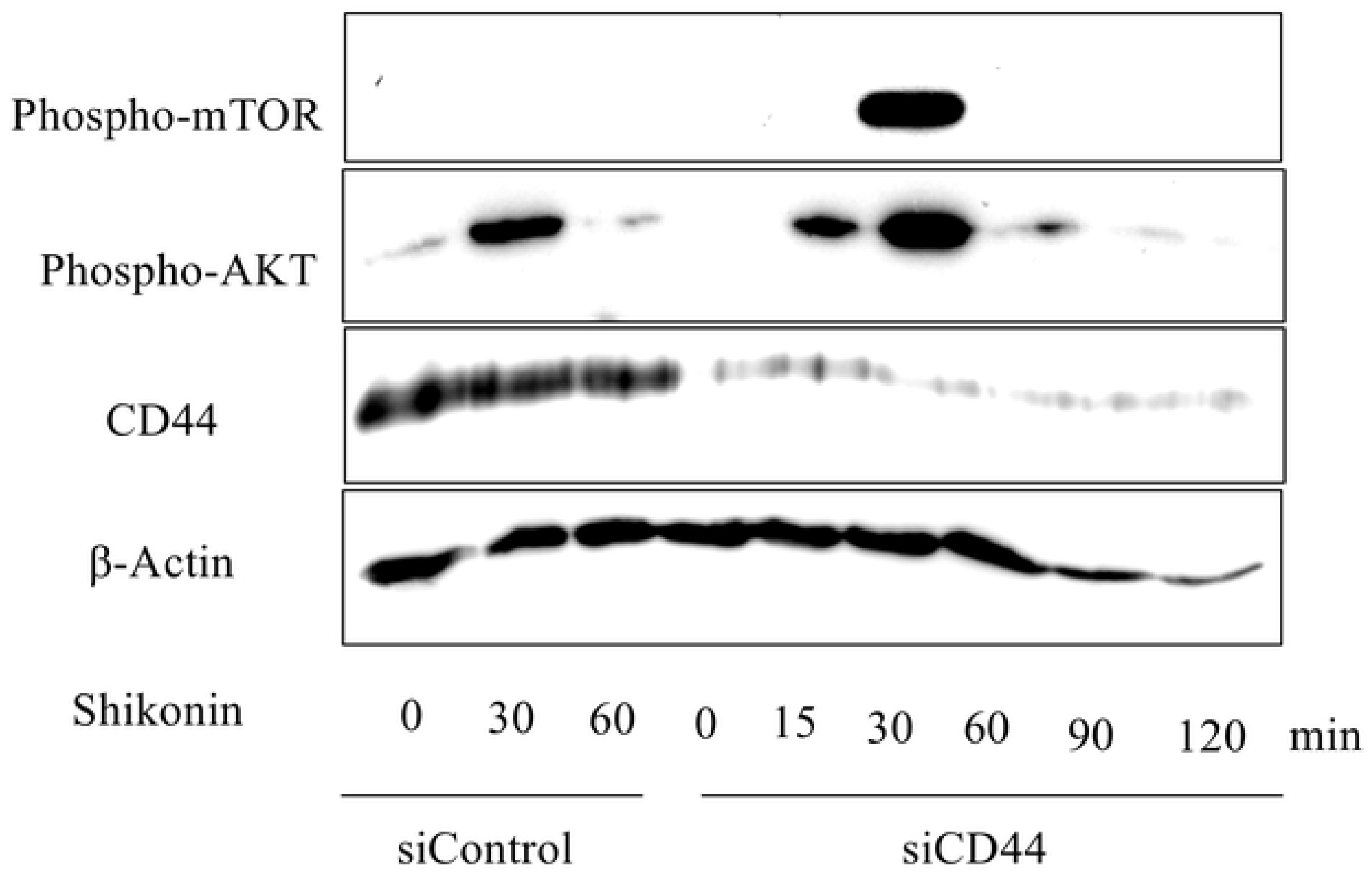
Shikonin Induces Expression of DSPP by a CD44-mediated Mechanism. DSPP expression levels after treatment with 0.5 μM shikonin in cells transfected with siRNA against CD44 (A). Changes in phospho-mTOR and phospho-AKT expression levels after treatment with 0.5 μM shikonin in cells transfected with siRNA against CD44 (B).

## Discussion

Our previous study indicated that hyaluronan induces odontoblastic differentiation of DPSCs via CD44 [1]. Because hyaluronan is a type of glycosaminoglycan with a molecular weight of up to 2000 kDa, it is necessary to explore novel small molecules that can induce odontoblastic differentiation for future clinical applications. Furthermore, a previous study revealed that CD44 expression was important for odontoblastic differentiation of DPSCs [1]. Hence, we performed this study to explore whether novel small molecules, not previously associated with CD44 expression, could induce odontoblastic differentiation of DPSCs. The aim of this study was to determine whether CD44 expression in DPSCs remained important for the activity of any novel compounds that could induce odontoblastic differentiation.

Vitamin K is a lipophilic vitamin involved in post-translational protein modifications associated with various blood coagulation factors [19]. Furthermore, it is involved in the production of vitamin K-dependent bone matrix proteins, such as osteocalcin [20]. Menatetrenone, a vitamin K analogue, has been used as an osteoporosis therapeutic agent or a hemostatic agent [21]. Therefore, we focused on vitamin K analogues in this study, and found that shikonin could induce odontoblastic differentiation. Shikonin is known as a natural red pigment component in herbal medicine, lithospermum root (Chinese name: Zi Cao). The main plant used for isolation of lithospermum root is *Lithospermum erythrorhizon*, a perennial plant native to East Asia with thick and purple roots that grows naturally in grasslands. The root is used as herbal medicine, and shikonin is the main component of lithospermum root (containing 0.5%–1.5%). In the second century ancient China, the drug book “Shennong Bencaojing” (from the later Han Dynasty to the Three States Period) lists lithospermum root as a medicinal herb; its medicinal properties are useful for the treatment of various diseases (e.g., pharyngitis, burns, cuts, measles, and purulent skin inflammation) [22]. Shikonin has been reported to inhibit growth of various cancer cells and induce apoptosis of these cells, as well as to suppress angiogenesis [23]. However, the underlying molecular mechanisms of these effects mediated by shikonin are not fully understood. Because we found that shikonin could induce odontoblastic differentiation of DPSCs in an enantioselective manner, we examined the intracellular signaling involved. The results showed that shikonin induced odontoblastic differentiation through CD44 and AKT–mTOR signaling. In addition, CD44 may be more critical for the shikonin-induced odontoblastic differentiation of DPSCs, compared with AKT–mTOR signaling, because CD44 inhibition suppressed odontoblastic differentiation regardless of changes in phosphorylation of AKT–mTOR. Thus, shikonin induces odontoblastic differentiation via AKT–mTOR signaling in the presence of CD44.

The PI3K/AKT pathway is a pro-proliferation signaling pathway that is active in various cells [24]. Furthermore, AKT is regarded as a positive regulator of the mTOR complex [25,26]. The AKT signaling pathway is reportedly involved in osteogenic differentiation of human mesenchymal stem cells, human dental follicle cells, and rat bone marrow stromal cells [27–29]. However, AKT reportedly inhibits odontogenic differentiation of stem cells from the apical papilla [30,31]; thus, there is controversy regarding the function of AKT–mTOR signaling. The results of our study suggest that the AKT–mTOR signaling pathway is involved in the shikonin-induced odontoblastic differentiation of DPSCs. Furthermore, CD44 was required for shikonin-induced odontoblastic differentiation.

Our study showed that shikonin could induce odontoblastic differentiation of DPSCs. The AKT–mTOR signaling pathway was the main intracellular mechanism involved; moreover, CD44 was important for the induction of odontoblastic differentiation. However, it remains unclear how the AKT–mTOR signaling pathway and CD44 are specifically related to the shikonin-induced odontoblastic differentiation of DPSCs. In addition, it remains unknown whether this effect of shikonin is present in vivo.

Our previous study indicated that the hyaluronan-induced odontoblastic differentiation of DPSCs was mediated by CD44. In the present study, the expression of CD44 was found to be important for inducing the odontoblastic differentiation of DPSCs, regardless of the presence or absence of treatment with hyaluronan. To the best of our knowledge, this study is the first report that shikonin can induce odontoblastic differentiation of DPSCs via CD44 and the AKT–mTOR signaling pathway. Because CD44-expressing cells are abundant in incomplete apical areas, pulp horn, and pulp chamber roof [15,16], shikonin may be a useful inducer of odontoblastic differentiation, and may have potential clinical use in the protection of pulp.

## Acknowledgments

We thank Ryan Chastain-Gross, Ph.D., from Edanz Group (https://en-author-services.edanzgroup.com) for editing a draft of this manuscript.

